# HTX: a tool for the exploration and visualization of high-throughput image assays

**DOI:** 10.1101/204016

**Authors:** Carlos Arteta, Victor Lempitsky, Jaroslav Zak, Xin Lu, J. Alison Noble, Andrew Zisserman

## Abstract

High-throughput screening (HTS) techniques have enabled large scale image-based studies, but extracting biological insights from the imaging data in an exploratory setting remains a challenge. Existing packages for this task either require expert annotations, which can bias the outcome of the study, or are completely unsupervised, failing to leverage the information present in the assay design. We present HTX, an interactive tool to aid in the exploration of large microscopy data sets by allowing the visualization of entire image-based assays according to visual similarities between the samples in an intuitive and navigable manner. Underlying HTX are a collection of novel algorithmic techniques for deep texture descriptor learning, 2D data visualization, adversarial suppression of batch effects, and backprop-based image saliency estimation.

We demonstrate that HTX can exploit the screen meta-data in order to learn screen-specific image descriptors, which are then used to quantify the visual similarity between samples in the assay. Given these similarities and the different visualization resources of HTX, it is shown that screens of small-molecule libraries on cell data can be easily explored, reproducing the results of previous studies where highly-specific domain knowledge was required.

## 1 Introduction

High-throughput screening (HTS) and high-content screening (HCS) techniques have enabled large scale image-based studies, where the effects of thousands of different treatments, such as small-molecules (e.g. Sigma Aldrich LOPAC or Pharmakon compound libraries) on one or more targets, are screened in a fully-automatic or semi-automatic way. Having automated ways of collecting the microscopy data translates into vast image collections being generated for each experiment, thus turning the data analysis into a major bottleneck in the experimental process. This has in turn motivated the development of automated analysis pipelines which have seen a rapid progress over the past decade, and more recently, due to the many successful efforts to adapt efficient and robust deep learning methods into microscopy image analysis. Nevertheless, extracting biological insights from large microscopy data sets in an *exploratory* setting remains a challenge.

Existing bioimaging packages based on image processing and machine learning have helped to alleviate the high-throughput analysis problem by providing the possibility of quantifying the relative effect of a compound, for example, by classifying individual cells into predefined phenotypes and comparing the phenotypic distribution of compound-treated versus control-treated cells. However, in the case of exploratory studies, the space of possible effects is unknown, thus quantifying only specific and specified effects can lead to important information being missed. Alternatively, methods based on unsupervised learning have the capability of recovering novel features from large image collections without direct human intervention, but they require good underlying visual features that can capture the patterns of relevance in each scenario. Moreover, due to the large amount of information that can be present in a large exploratory screen, interactive elements in the data visualization can help to improve the workflow by providing flexibility to query the data, enabling the identification of novel features.

We present HTX (High-Throughput eXplorer), an interactive tool to aid in the exploration of large microscopy data sets by allowing the visualization of entire image-based assays according to visual similarities between the samples in an intuitive and navigable manner. As an exploratory tool, HTX can be used to complement traditional High-Content Screening software. HTX benefits from generating image descriptors for any target data set through the use of deep learning with convolutional networks using only the metadata provided with the high-throughput assay. Furthermore, given the possibility of quantifying the similarity between biological samples, HTX leverages methods for image saliency in deep networks to explore the characteristic visual patterns that arise in a screen.

Aside from proposing a novel pipeline for the exploration of large microscopy screens, within HTX we make the following technical contributions:

- We propose a new way to learn deep texture descriptors for high-throughput screens without human-provided image labels.
- We show that the usage of adversarial training in deep neural networks allows to reduce batch effects in the analysis of HTS data sets.
- We propose a diffusion process-based approach for creating a diverse family of 2D visualizations at multiple levels of detail.
- Finally, we propose an approach for on-the-fly image saliency computation from deep neural networks that can be controlled interactively by the user.

The different visualization resources of HTX are demonstrated on two publicly available high-throughput screenings of small-molecule libraries on cell data: the BBBC022v1 data set [11], and BBBC021v1 [2], both from the Broad Bioimage Benchmark Collection [17].

## 2 Related work

Existing methods for the exploration of high-throughput microscopy data sets can be broadly grouped into two main approaches, those which rely on some form of direct human supervision at the image level, and those which rely on unsupervised learning algorithms. Broader reviews of the available techniques can be found in [10, 15].

### Supervised classification approaches

A common approach for parsing large data sets of microscopy images is by learning phenotype classifiers of regions of interest (e.g. cell segmentation), and comparing the phenotypic distribution of treated versus control samples. Traditionally, these classification pipelines would be based on hand-crafted visual features (e.g. cell texture or morphology descriptors) and learning algorithms such as SVMs and K-NN, as available in popular open software packages [5, 13, 14] or software integrated in high-content imaging systems. More recent methods in this category follow the trends in machine learning and make use of deep hierarchical feature architectures to learn the phenotype classifiers end-to-end [9, 16, 18, 24]. The classification-based approach is expected to work very well when applicable. For example, deep learning architectures have been shown to be very powerful tools for solving classification problems in a wide variety of scenarios, including biomedical imaging. However, in practice, casting the microscopy data set exploration as a classification task requires the analyst to first decide what the classes of interest are (e.g. a discrete set of phenotypes), find a set of examples for each of the classes, and then train the network to recognize those classes. Instead, if one is concerned with exploring the screen without introducing discrete classes, due to impossibility or with the intention of discovering new features in the data in an unbiased manner, then the classification approach is not directly applicable.

### Unsupervised approaches

Approaches based on unsupervised learning algorithms tend to be more applicable to exploratory scenarios where prior information about the range of visual patterns in the data is unavailable. Within this category we find clustering-based methods [11, 12, 19, 29], as well as methods based on dimensionality reduction, usually with the end goal of data set visualization [21, 22]. With sufficient domain knowledge, unsupervised methods can be used to visualize, explore and discover novel features in large microscopy data sets, but it is often difficult to assess the significance of the results [20], and additional exploration and visualization resources are required. Moreover, similar to the classification approach, the methods in this category build on top of visual features that are commonly hand-crafted, and thus have to be chosen in advance according to prior knowledge about the experiment. It is also possible to learn features specific for the target data, but unlike the classification approach, making use of novel deep learning architecture has not yet become as popular for unsupervised methods given that feature learning with little or no supervision is a more challenging learning problem. Our approach leverages the information in the screen metadata as a form of supervision in order to learn screen-specific image descriptors without the need for human labeling.

## 3 Methods

### 3.1 Learning texture descriptors

The core of HTX is the capability of quantifying general visual similarities between microscopy images in an unbiased manner; that is, without the necessity of predefining the specific visual patterns that should be taken into account. This is achieved through the application of modern computer vision techniques for the measurement of texture similarities based on descriptors learned from data. Instead of using hand-engineered image features, as commonly done in machine learning pipelines for microscopy image analysis, including similar exploratory tools, HTX makes use of end-to-end learnable *deep texture descriptors* [8]. These texture descriptors are based on Gram matrices computed from activation maps in a deep convolutional network as shown in **Fig. 2**. Each coefficient of a Gram matrix corresponds to a spatially-pooled correlation between various pairs of convolutional feature maps. Since the feature maps are progressively down-sampled through the CNN, the three Gram matrices capture textures information at different spatial scales.

The texture descriptors in HTX are initially learned on a data set of natural textures through a *pre-training* process (**Fig. 1a**) done to the encoding network in a data set for texture classification of textures in the wild [3], as well as synthetic textures [28]. Once initialized, the descriptors in the texture classification network are adapted to any specific HTS data set through a simple *fine-tuning step* (**Fig. 1b**). Fine-tuning the texture description network only requires the metadata associated with the screen as a form of supervision, as opposed to, for example, labeling the data by phenotype, which can be labour-intensive or subjective. For instance, HTX can make use of the information about sister wells (i.e. wells treated with the same compound, for which it is desirable to have similar textural descriptors) to establish similarity relations between samples, and use those to fine-tune the parameters of the texture-extracting convolutional network. The similarity relations between samples are enforced during learning through the histogram loss [25] (**Fig. 3**), resulting in an image embedding specific for the target screen.

**Figure 1:**
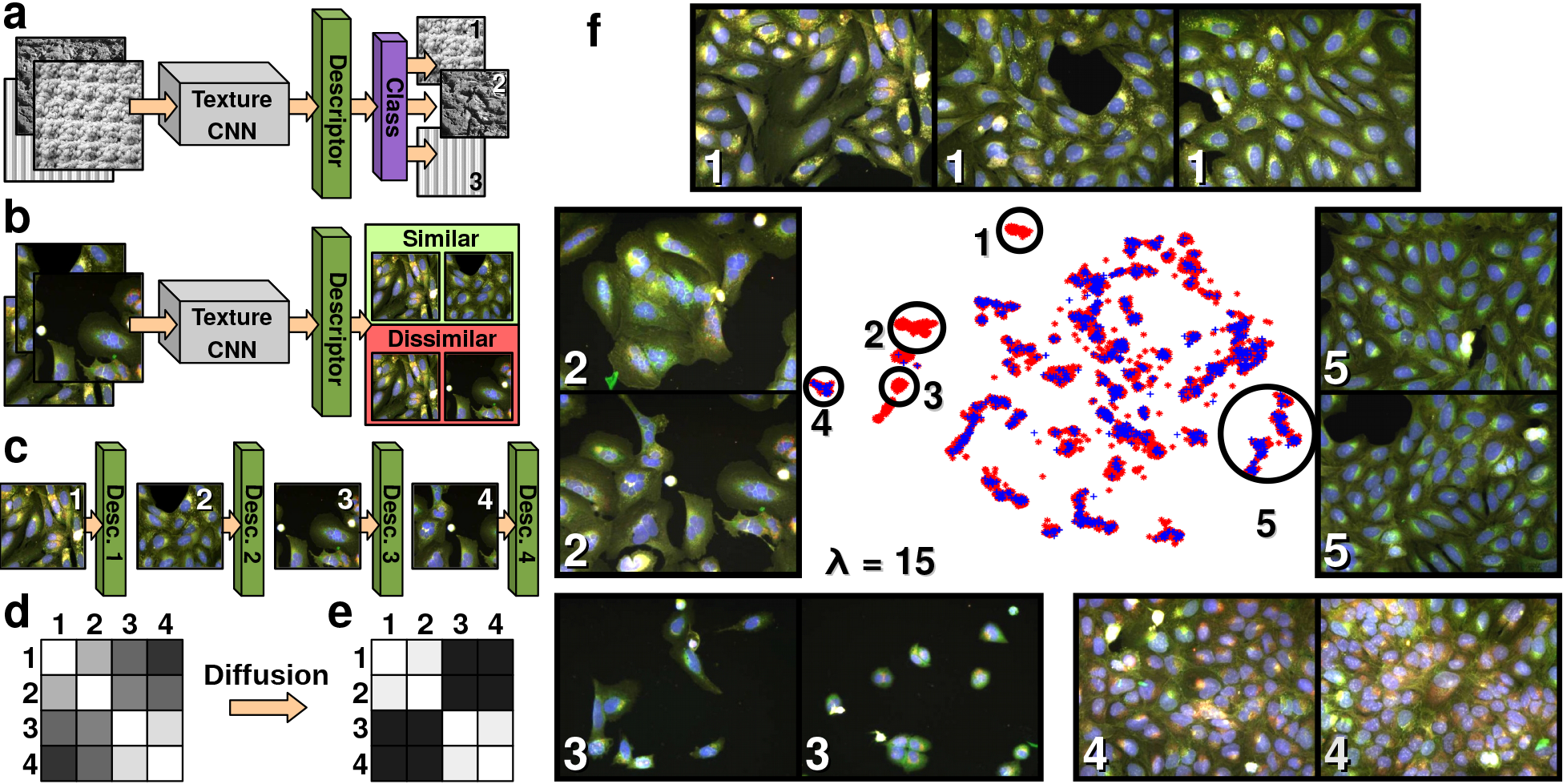
Visualizing a high-throughput screening according to the visual similarities in its samples. **(a)** The core of HTX is a Convolutional Neural Network (CNN) that is trained to recognize and describe image textures. **(b)** Given a target HTS data set, the texture encoding CNN is fine-tuned by learning similarity and dissimilarity relations between the samples in the data set, along with a special mechanism to reduce the effect of batches in the screen, shown in **Fig. 3**. **(c)** Once the CNN is adapted to the HTS domain, it can encode all the samples in the data set. **(d)** The sample descriptors can then be used to measure the affinity between any pair of samples, resulting in an *affinity matrix* (e.g. **Supplementary Fig. 1**). **(e)** The similarities in the affinity matrix can be diffused to emphasize local or global structures. **(f)** From the affinity matrix, all samples in the HTS data set (red asterisks for the compounds being explored, and blue crosses for control samples) are projected into a 2-D space where samples lying near to each other are considered visually similar. Therefore, exploring the different naturally-formed clusters in the 2-D space gives an idea of the different range of visual effects in the experiment. Clusters with little or no presence of control samples (e.g. 1, 2, & 3) can be a quick indicative of active compounds.

Due to the plate-based operation of HTS systems, along with minor imperfections, HTS data sets often contain batch effects which are undesirable as they can bias the similarity measures between samples; i.e., samples in the same plate or batch can appear more similar between them than to those in separate batches, regardless of the experimental variables. In order to alleviate this effect, the encoding CNN is trained with an auxiliary mechanism that drives the network to *not* learn the distinctions between different plates. This reduction of batch effects is implemented through adversarial domain adaptation [7], which is shown in **Fig. 3**. Intuitively, a network classification branch is trained to predict from which plate a sample is coming, while an additional adversarial branch attempts to affect the CNN such that the plate classifier fails at its task. The plate classifier is implemented as a multiclass softmax classifier, with as many classes as plates in the data set, while the adversarial branch measures the KL-divergence between the output of the softmax and a uniform distribution, as suggested by Tzeng *et. al* [23]. The error gradients of the classification branch are only backpropagated through the plate classification network, whereas the gradients of the adversarial loss are back-propagated all the way through the encoding network, and these two objectives are alternated during the network training. Updating the encoding network such that the output of the classification network is a uniform distribution, as attempted by the adversarial branch, would be equivalent to making the encoding network ignore the visual feature that characterize the HTS plates.

### 3.2 Sample description and similarity

Once the texture description network is adapted for a target screen, it can then represent each sample in the screen with a descriptor *x_i_*, a vector that encodes its textural patterns (**Fig. 1c**). The sample descriptor is made of the concatenation of the vectorized Gram matrices computed from the activations of layers at different depths within a convolutional neural network (CNN), which is shown in full in **Fig. 2**. The *texture descriptor x_i_* can then be used to determine how visually similar two samples *i* and *j* are (e.g. by evaluating the distance ∥*x_i_* - *x_j_*∥), and to build a matrix of *pairwise affinities A* (**Fig. 1d**) for the entire data set (**Supplementary Fig. 1**).

**Figure 2:**
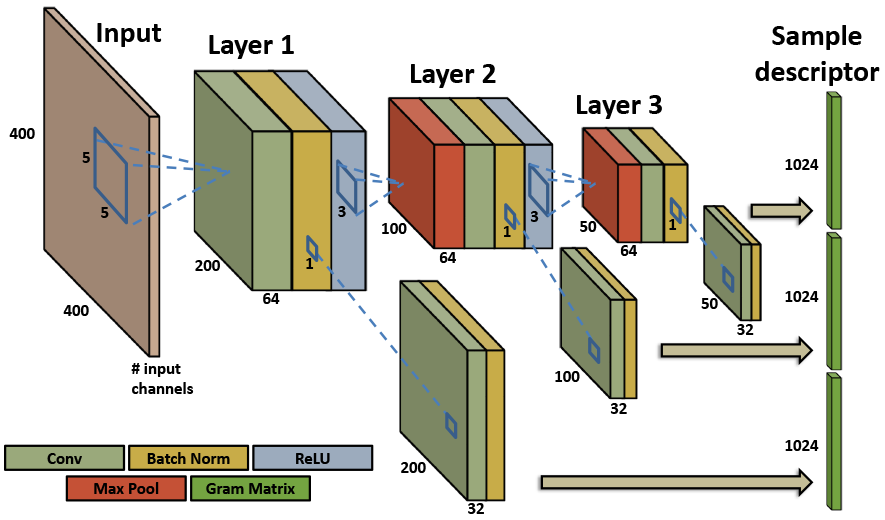
Architecture of the sample encoding CNN. Given an input image stack, the encoding CNN produces an encoding vector consisting on the concatenation of texture descriptors (Gram matrices) computed over the visual features a different depths of the network. *Conv, Batch Norm* and *ReLU* correspond to convolutional, batch normalization and rectified linear unit layers.

### 3.3 Multi-level visualizations

Given the matrix *A* containing the affinities between every pair of samples in the data set, HTX can present all the relations within it to the user in an intuitive way, by projecting the samples into a two-dimensional space that can be easily interpreted and managed (**Fig. 1f**). A low-dimensional projection of high-dimensional data generally comes with a compromise on the accuracy of the representation; in order to alleviate this problem, HTX makes use of the *t-distributed stochastic embedding* (t-SNE) method [26], a nonlinear dimensionality reduction method specially developed for the visualization of high-dimensional data. t-SNE begins by building the distribution *P*, containing the conditional probabilities *P_i|j_* reflecting the pairwise affinities between the highdimensional vectors representing the samples (*A_i,j_*). An analogous distribution *Q* is then built for the samples 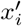 in the lower-dimensional space, and finally, the values of 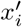 are searched such that the Kullback-Leibler Divergence between the distributions *P* and *Q* is minimized.

T-SNE is capable of capturing the structures in the manifold where the high-dimensional image descriptors lie. Therefore, by emphasizing local or global structures in the manifold, it is possible to obtain visualizations of the entire data set at different levels of granularity. The level of granularity of the t-SNE visualization can be changed by smoothing the affinities in A using *diffusion* (**Fig. 1e**) [6] on the high-dimensional manifold. Generally, stronger smoothing leads to less fragmented manifold visualizations. In practice, the propagation is done on an affinity graph built from *A*, where every sample *s_i_* is a node, and the weights between nodes *s_i_* and *s_j_* is given by their affinity *A_i,j_*. The affinities are then diffused from each node to its *k*-nearest neighbours, and thus, *k* controls the extent of the diffusion process.

HTX computes the t-SNE 2-D visualization over affinity matrices for various extents of diffusion, generating visualizations at different levels of detail that the user can navigate through (**Fig. 4a**).

### 3.4 Highlighting salient visual features

The 2-D visualizations (e.g. (**Fig. 1f**) are generally populated with naturally-formed clusters of visually similar images. Nevertheless, it is not always obvious what thespecific patterns are that make samples in a cluster similar to each other and dissimilar to other images (c.f. samples from same clusters in **Fig. 1f**). Likewise, it is possible that the visual patterns that the network consider to be particular to a cluster differ from what a user can visually interpret. To address this possible visual ambiguity, HTX has the capability of *highlighting* the image regions that most contribute to discriminating images in the cluster from others. For each image in a cluster *C_k_*, the highlighting procedure creates a saliency map that displays the relative contribution of every image region towards the formation of the cluster (**Fig. 4b**) by determining how important every local feature is to discriminate *C_K_* from other clusters. Since our descriptors are based on deep neural network, we can exploit visualization algorithms based on back-propagated error signals. For this purpose, we make use of the excitation backpropagation (EBP) method [30].

Class saliency methods (such as EPB) are usually applied to obtain saliency masks for the classes observed during training. However, in our setting, not only the network is not trained for classification, but as a data exploration scenario, the groups of samples to be compared are defined as part of an interactive process. As the user selects a certain cluster of interest in the t-SNE visualization, the saliency maps indicating spatial locations and channels that distinguish the selected cluster from the remainder of the data set must be computed on-the-fly for the members of that cluster. Therefore, EBP is only applicable to our encoding CNN after the following required modification. Given the user-defined classes corresponding to sets of images specified (circled) by the user on the t-SNE visualization (e.g. a cluster of treated samples in the t-SNE visualization and a reference group to compare them against such as all control samples), we train a linear SVM to discriminate between the two groups using the sample descriptors as training samples, which are loaded from memory. The weights of the SVM classifier are then transplanted into a fully-connected classification layer that turns our encoding CNN into a binary classifier for the two classes (the selected cluster and the reference group). Such a procedure allows for any of the common class saliency methods for deep architectures to be used for the selected groups, including our choice of EBP. We compute class saliency for the input of the network, thus obtaining as many saliency maps as there are input image channels. Finally, these saliency maps are blended into RGB channels to form the images in **Fig. 4b**. These saliency maps can then be displayed alongside the original compound images as the user browses the members of the cluster. The interactive exploration process is demonstrated in **Supplementary Video 1**.

### 3.5 Measuring cluster enrichments

In exploratory tools for HTS, it is also important to provide the means to quantitatively validate hypotheses that arise during the data set exploration. A way of doing so that is common in the analysis of small-molecule screens or similar high-throughput assays, is the computation of enrichments of treatment properties (e.g. therapeutic use of the small-molecule) for a particular data subset arising in the exploration. For example, a subset of interest can be the a cluster of samples that appear in our data set visualizations (e.g. **Fig. 1f**). Therefore, the HTX interface allows not only to visualize the data set and the salient visual patterns in the naturally-formed clusters, but also provides a simple way to evaluate enrichments (e.g. using the screen metadata) on interesting clusters in order to establish hypotheses that are quantitatively supported. Note that the definition of clusters can be done not only interactively by the user, but also automatically. In this *offline* modality, soft-clusters, or neighbourhoods 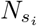 around each sample *s_i_* can be defined directly from the affinities in the matrix *A*, and the enrichments can be pre-computed to provide to the user a set of potentially active neighbourhoods. Such neighbourhoods can also be visualized in the 2D maps, as shown in **Supplementary Video 2**, and explored with the visual saliency tool.

### 3.6 Implementation details

#### Network training details

The entire network in **Fig. 3**, consisting of the encoding network plus the batch effect reduction branch, is trained simultaneously and end-to-end using the backpropagation-based stochastic gradient descent with with momentum. The images are preprocessed by subtracting a global channel mean, computed over the entire data set; no other pre-processing is applied. The training data is augmented by taking random crops and rotations of the input images before passing them through the network. During pre-training on the texture classification datasets [3, 28], the input of the network is single-channel (grayscale) texture images. During the fine-tuning on the data set of interest, the first convolutional layer is replicated as many times as there are input channels in the data set. For example, if the original convolution tensor in the first convolutional layer is of size *ks × ks × 1 × outSize* during pre-training, the tensor gets replicated to *ks × ks × 5 × outSize* in order to handle the 5 input channels of the BBBC022v1 data set, where *ks* is the spatial kernel size in the spatial dimension. The network implementation and training was done using the MatConvNet toolbox [27]. Learning the embedding for the BBBC022v1 data set takes approximately 2 days on a Titan-X GPU; this duration is largely dominated by the heavy reading operations from disk of TIFF stacks with 5-channels, 520x696 pixels and 16-bit resolution.

**Figure 3:**
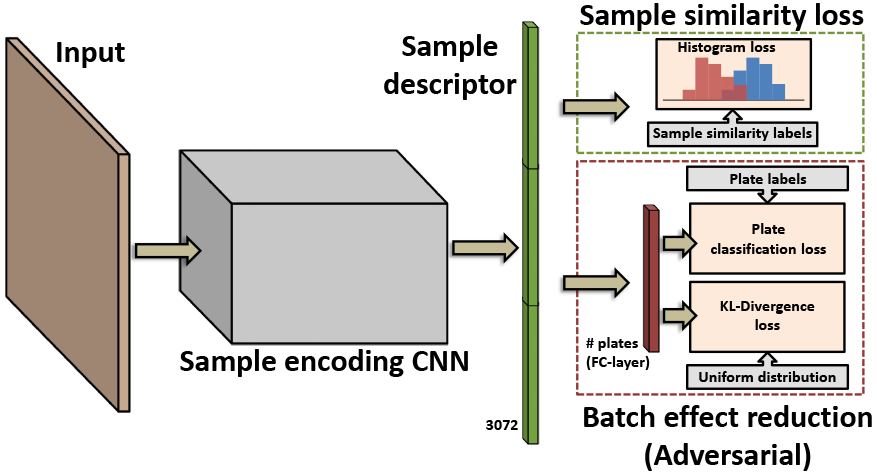
Training the sample encoding CNN. The sample encoding CNN, which is initially trained to recognize image textures, is adapted to a target HTS data set through a multi-task training procedure. The core loss function, the *sample similarity loss*, takes groups of samples along with information about their relation (i.e. similar or dissimilar) and tries to minimize the probability that the similarity between dissimilar samples would be higher than the similarity between similar samples. Simultaneously, a *batch effect reduction* branch attempts to prevent the network from using batch/plate-specific visual features to solve the similarity task.

#### Discarding empty images during network training

In wells of low confluency, it is possible to obtain some empty fields-of-view when randomly sampling a well. This is undesirable when learning the similarity measures, since an empty frame could be labeled as similar to a nonempty frame by the metadata, forcing the network to assimilate them as such. Nevertheless, this potential issues can be easily avoided with any cell detection approach. In the case of the data sets used in this paper, it is straightforward to obtain nuclei and non-nuclei regions by applying a simple automatic thresholding (Otsu’s method) operation on the nuclei channel of the images within a well. Following this thresholding, empty images are discarded if they do not contain any cell nuclei.

#### Automatic neighbourhood definition

When computing the enrichment of some property (e.g. compound-concentration pairs) on groups of samples, as explained below, we make a distinction between clusters in the 2-D t-SNE visualization, and the neighbourhoods in the high dimensional texture descriptor space. Indeed, projecting relative distances of high-dimensional vectors onto a 2D space will incur severe distortions. Therefore, when automatically defining the samples that lie next to some sample *s_i_* (i.e. the neighbourhood 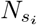), we consider the likely neighbours of *s_i_* in the high-dimensional descriptor space, and not in the 2-D visualization. To achieve this, we follow the approach taken by the t-SNE visualization technique, and place a Gaussian kernel *G_i_* centered at each sample *s_i_*, which defines a distribution over all other samples *s_j_* that indicates how likely they are to be the neighbour of *s_i_*. The variance *σ_i_* of the Gaussian kernel would be required to change according to the local sample density in the high-dimensional space. Therefore, instead of defining *σ_i_*, a global parameter *P* is set corresponding to the desired perplexity of the Gaussian kernels *G*, which in turns adapts *σ_i_* according to the neighbourhood of *s_i_*. Once the distribution around each sample *s_i_* is obtained, the neighbourhood 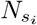 is defined as the samples within 0.99 probability of *G_i_*. We use a perplexity *P* = 5.

#### Measuring enrichments

In the data set analysis examples presented in the paper, we show the computation of the compound enrichment for user-selected clusters from the 2-D visualizations, or sample neighbourhoods that are automatically defined. In either case, the enrichment is evaluated for each compound-concentration pair in the database using the hypergeometric test with Bonferroni correction, for *p* ≤ 0.01.

## 4 Example results and discussion

We now demonstrate the applicability of the tools in HTX, and how they facilitate exploring real HTS data while leading to the same conclusions as specialized pipelines for cell image analysis. We use HTX to visualize two publicly available data sets of cell-based HTS from the Broad Bioimage Benchmark Collection [17]. Note that direct comparisons of exploratory tools on real biological data based on accuracy is often not possible given the subjectivity of the task, especially when interactive elements are present, as it is common in this setting.

When fine-tuning the texture description network in each of the two examples screens, the similarity-dissimilarity relations between samples are set in the following way: two samples are considered *similar* if they have been treated with the same compound and concentration, and *dissimilar* otherwise. Additionally, during fine-tuning, the multiple images from a single well are treated separately to maximize the amount of training samples. However, when producing the final descriptor *x_i_* for well *s_i_*, we average the descriptor of each of the images in it. Finally, the similarity between two samples *s_i_* and *s_j_* is measured by taking the euclidean distance between their descriptors (‖*x_i_* − *x_j_* ‖) following L2-normalization.

### 4.1 BBBC022v1 screen

First, we use HTX to explore the BBBC022v1 data set [11] (**Figs. 1** and **2**), which consists of a screen of 1600 known bioactive compounds (with biological replicates) on the osteosarcoma cell line U2OS. The wells are imaged at 9 different fields of view and 5 channels, each containing different cellular components: nucleus, nucleoli, Golgi apparatus and plasma membrane, F-actin, and mitochondria. The screen contained 20 plates of 384 wells, for a total of 7,680 samples, all of which are shown in the 2-D visualizations of (**Fig. 1f**). Here, blue crosses correspond to the control samples and the red asterisk to the treated wells. A key observation in the 2-D maps is that, at all levels of granularity (e.g. **Fig. 4a**), clusters composed of test compounds devoid of any control samples can be identified. This demonstrates the ability of HTX to distinguish active from inactive compounds within the assay. The clusters of active compounds identified in the 2-D maps (**Fig. 4a**), shown in **Fig. 4b**, were selected for further inspection and were found to overlap with the clusters of active compounds discovered by Gustafsdottir *et. al* [11] through a pipeline specific for cell images, thus confirming their relevance. We then used the saliency tool to assess the distinctive visual patterns of each of these cluster. As shown in **Fig. 4b**, the saliency tool is able to highlightthe features that most distinctively characterize each of the three active clusters. The most strongly highlighted patterns corresponded to a pronounced Golgi staining in cluster 1, multi-nucleation in cluster 2, and high cytotoxicity and plasma membrane blebbing in cluster 3, thus giving an indication of the range of effects that can be captured with the learned texture descriptors. Finally, these clusters with active compounds were also identified automatically through enrichments from the assay metadata (**Supplementary Video 2**). The enrichments were computed based on the annotation terms provided by Gustafsdottir *et. al* [11] for a subset of the compounds, which come from sources such as Gene Ontology [4], medical subject headings (MeSH) and the use/class of the compounds. As shown in **Supplementary Video 2**, the top enrichments for such annotation terms were found, as expected, within the active clusters.

**Figure 4:**
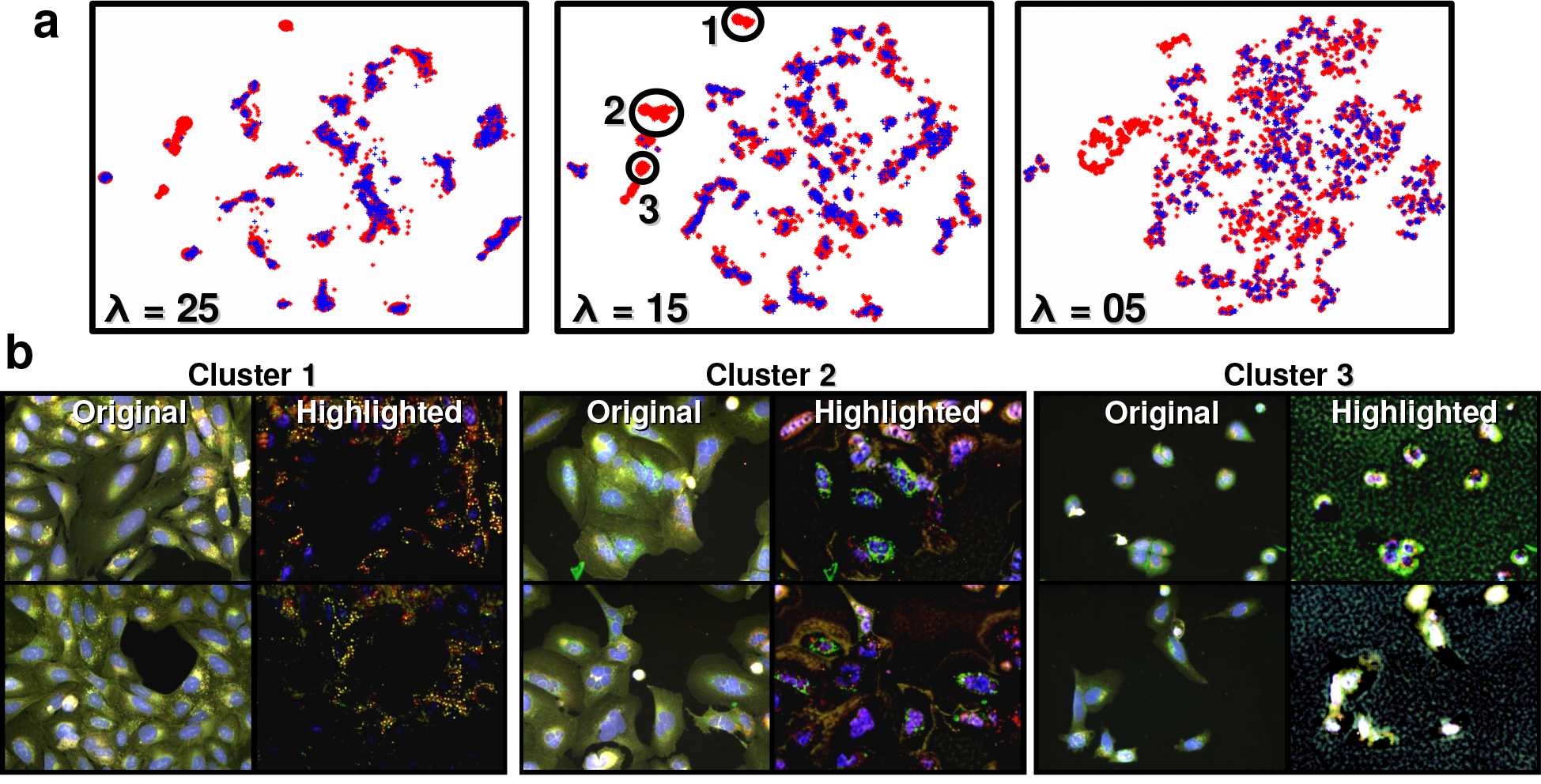
Interacting with the 2-D visualizations. **(a)** The level of detail considered in the 2-D visualizations (**Fig. 1**) can be varied with a parameter λ in HTX, which controls the level of similarity diffusion, resulting in finer or coarser visualizations of the data set. **(b)** Clusters in the 2-D visualizations can be selected in order to highlight the visual patterns that make the samples in the cluster similar (and different from the rest). The pattern highlighting feature allows for an objective judgement of the visual effects present in a cluster, as well as their relative importance. We show the distinctive pattern highlighting for three clusters selected from the 2-D visualization in **Fig. 1**; these cluster were chosen as they overlap with the active clusters found and described in the original BBBC022v1 data set analysis [11]. Cluster 1 mainly contains a pronounced Golgi staining. Cluster 2 contains multi-nucleated cells. Cluster 3 shows reduced cell size, condensed nuclei, plasma membrane blebbing, reduced nucleolar staining, and significant cytotoxicity. Such distinctive features automatically appear highlighted in our visualization.

Note that the multi-scale nature of the texture descriptor, in combination with its learnable design, allows to avoid common biases such as prioritizing image-level or global patterns (e.g. cell density) over more subtle local patterns, as shown in **Supplementary Fig. 2.**

### 4.2 BBBC021v1 screen

The second example data set is BBBC021v1, extracted from [2], and consists of a compound profiling assay on the MCF-7 breast cancer cell-line treated with 113 small molecules at eight different concentrations. The wells are imaged at 4 different fields of view and 3 channels: DNA, F-actin, and tubulin. A similar output to the exploration of BBBC022v1 is shown in **Fig. 5** and **Fig. 6**, where it is also visible, at all levels of granularity, naturally-formed clusters of samples (red asterisk) that do not contain control samples (blue crosses). As expected, these particular clusters reveal the most active compounds in the screen, and according to their relative distance in the visualization, also contain information about the similarities in their mechanism of actions [2].

**Figure 5:**
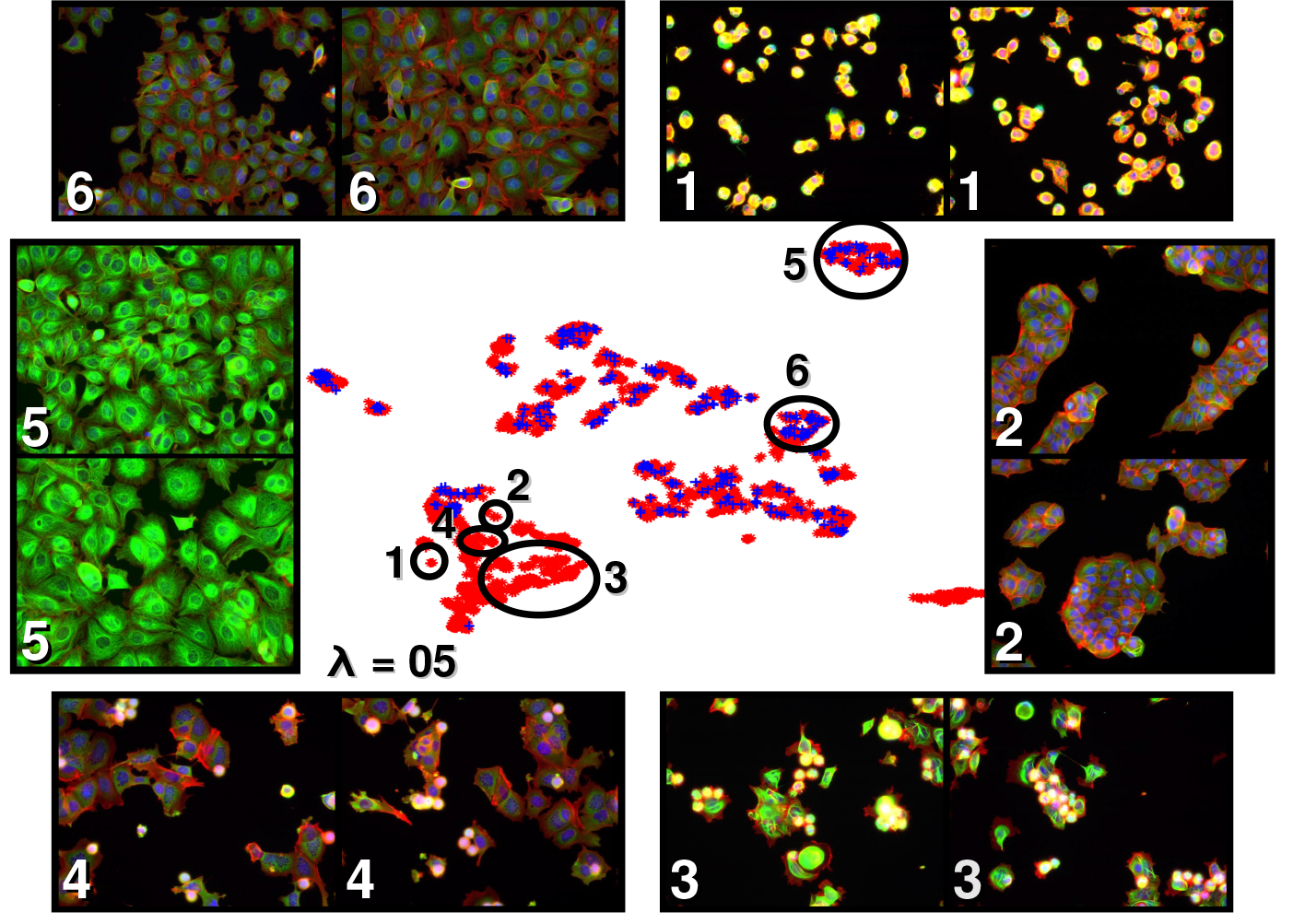
Visualization of the BBBC021v1 data set. The 2-D map in the centre is a projection of the entire BBBC021v1 data set, where red asterisks represent the compounds being explored, and blue crosses the control samples. Samples lying nearby to each other in the 2-D map are being considered more visually similar than those lying further apart. It is useful to note that some areas in the 2-D map are free from control samples, which is an indicative of biologically active compounds such as those in clusters 1-4. Likewise, samples within clusters containing controls are likely to contain inactive compounds (e.g. cluster 6). Additionally, clusters with control samples but appearing isolated, can correspond to defective samples (e.g. cluster 5).

**Figure 6:**
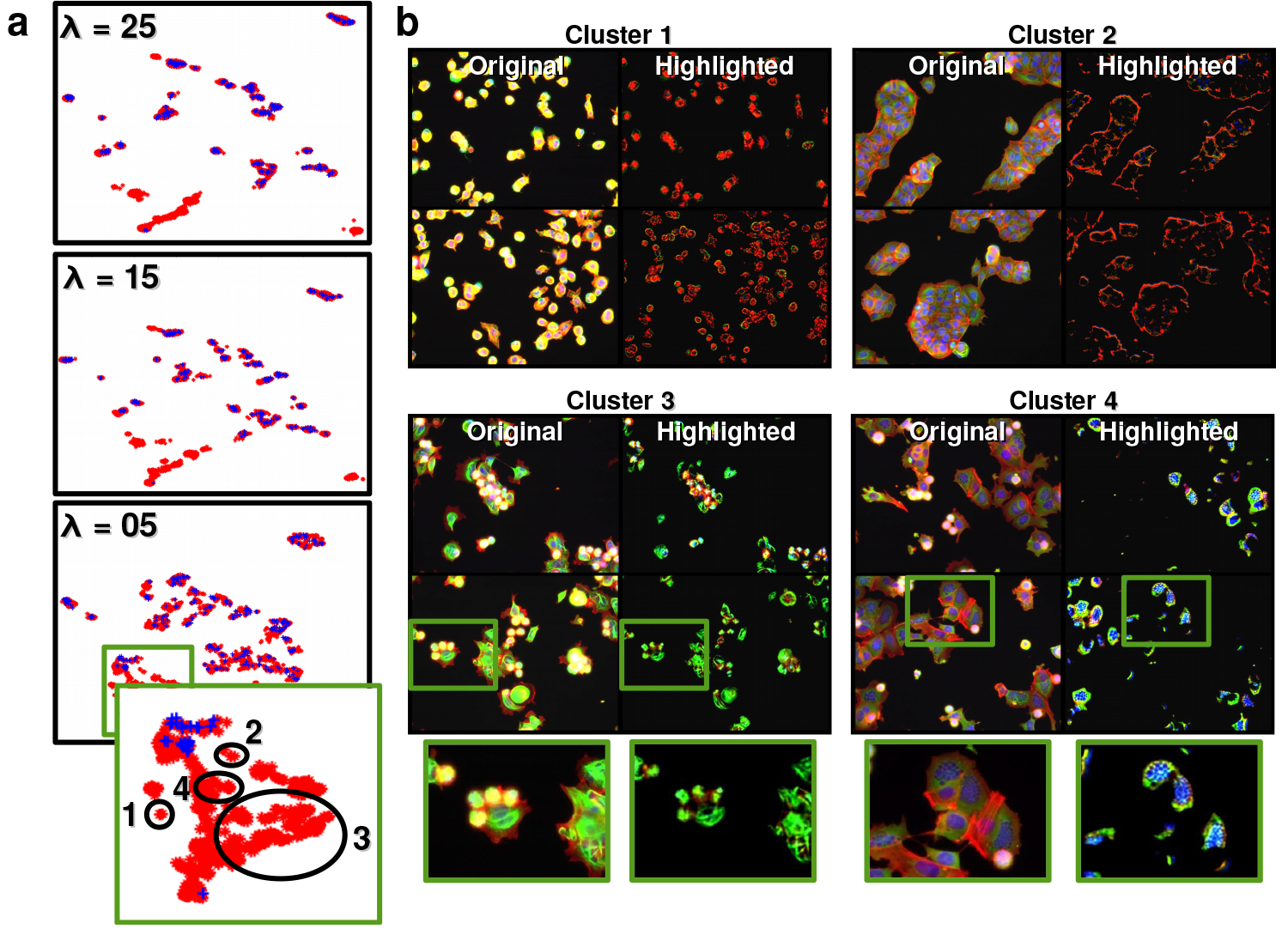
Interacting with the 2-D visualizations for the BBBC021v1 data set. **(a)** The level of detail considered in the 2-D visualizations (**Fig. 5**) can be varied with a parameter λ in HTX, which controls the level of similarity diffusion, resulting in finer of coarser visualizations of the data set. **(b)** Using the tool for highlighting relevant visual features in HTX, we visualize the salient patterns in clusters 1-4 from **Fig. 5**, located in the area of active samples on the 2-D map. Within these clusters, we find two active clusters that were also identified in a previous BBBC021v1 analysis [2]: cluster 3, containing compounds such as Taxol and Docetaxel, which act as microtubule destabilizers, and cluster 4, containing compounds such as Demecolcine and Vincristine, also microtubule destabilizers which additionally cause nuclear fragmentation.

This data set can be explored in the demo provided along with the source code of the tool [1].

## 5 Conclusion

We have introduced HTX, a computer vision and machine learning pipeline designed to be an aid in the visualization and exploration of high-throughput screens. Through the use of the different visualization resources in HTX, we have demonstrated how it can be used to explore real small-molecule screens, easily reaching similar conclusions (e.g. identification of active compounds) to those obtained through the use of setting-specific image analysis pipelines, and without the need of any human annotation of images.

Learning the texture descriptors specific for a new high-throughput data set can be a relatively expensive process computationally, with the main bottleneck being the loading of continuous streams of raw images, at the pixel depth and resolution commonly used by high-content screening systems. Nevertheless, this could be greatly alleviated with careful data preprocessing, which is not explored in this paper. Once the learning is performed, the data exploration is an interactive process where further computations, such as the feature highlighting tool, are done on-the-fly.

To conclude, we note that HTX is a general tool that can be applied to experimental setups other than cell imaging with fluorescence microscopy, including other microscopy modalities, organisms, or even visual data outside the microscopy domain. Such flexibility mainly arises from learning the image descriptors from the target data instead of manually selecting them for the specific domain.

## Acknowledgement

We thank Heba Sailem, Jens Rittscher and Anne Carpenter for very helpful discussions.

---

**Supplementary Fig. 1:**
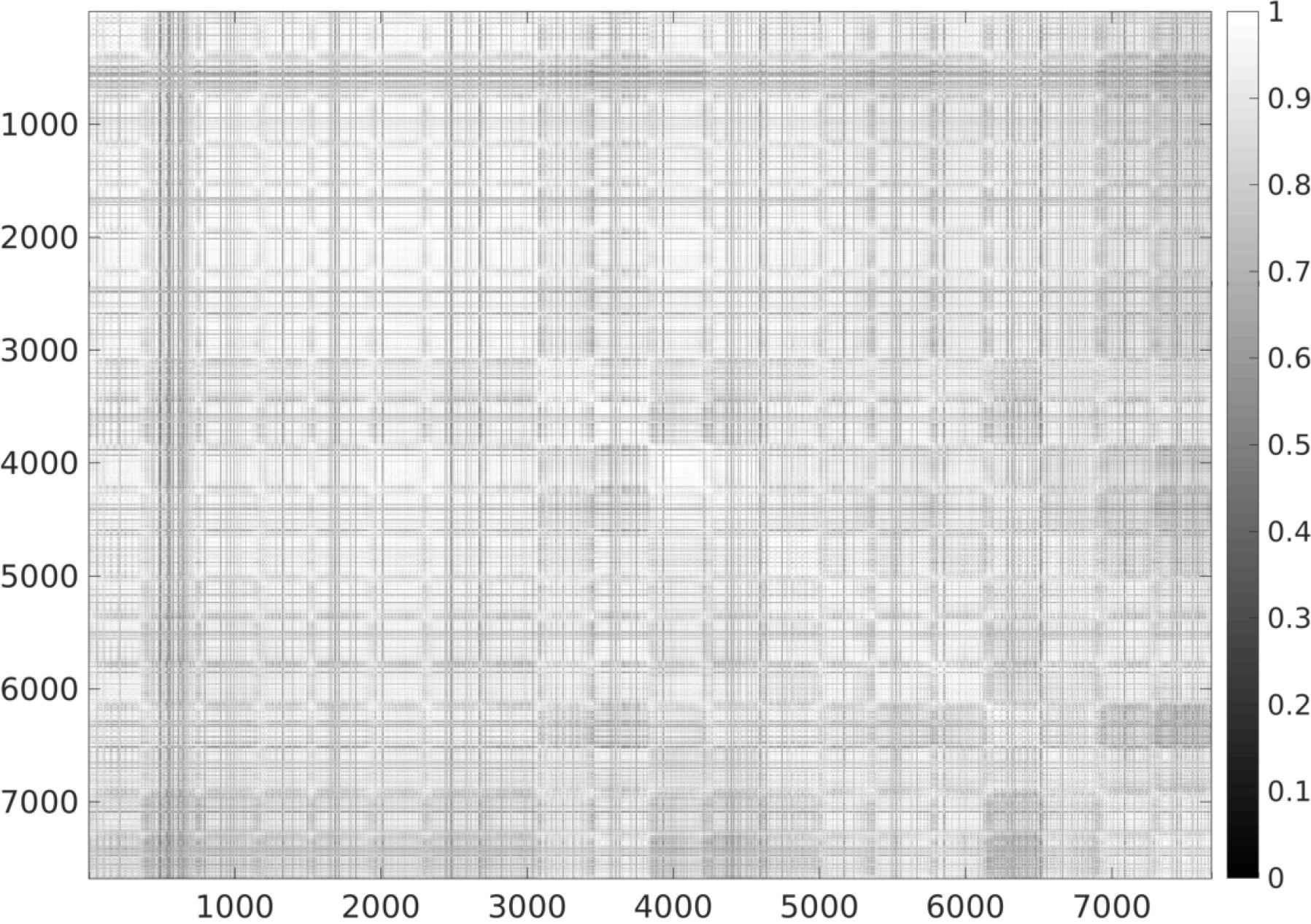
Affinity matrix for the 7681 samples of the BBBC022v1 data set.

**Supplementary Fig. 2:**
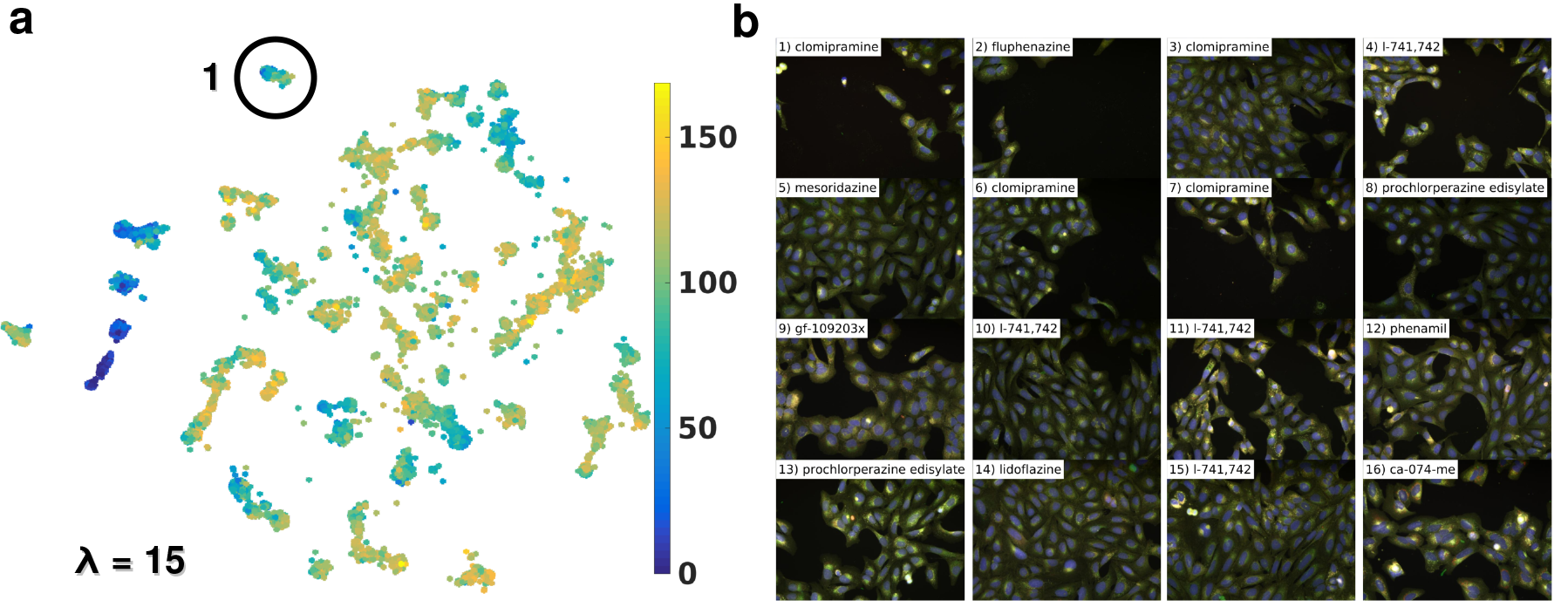
Effect of cell density on the sample similarity measures. In order to observe the effect of the patterns related to cell density, **(a)** we visualize the 2-D map of **Fig. 1f**, encoding in the color of each sample (i.e. an asterisk in the plot) the average number of cells per field-of-view. We can observe that the samples in the 2-D map are not clustered based purely on cell density, as indicated by the presence of several different clusters with similar cell densities, as well as clusters containing samples with varying cell densities. Most notably, cluster 1 contains a relevant phenotype mainly reflected in the pronounced Golgi staining, as described in the main text, along with relatively strong variations in cell density. To explore this example, **(b)** 16 samples from cluster 1 are shown. These samples were selected by choosing a sample with low cell density from cluster 1 (top left), and retrieving its most similar 15 samples in terms of the texture descriptors. Among those, we can observe significant variability in cell density, showing that the local Golgi staining was able to dominate the similarity measurement over the global cell density patterns. When measuring the similarity between samples in cell experiments, it is important that the computation is not biased by the strong patterns caused by variations in cell density. Although cell density could be relevant in many cases, and can appear correlated to other phenotypes (e.g. from toxic compounds), it has the potential to dominate over more subtle but more interesting phenotypes. The different scales of the texture descriptor in HTX range from receptive fields smaller than a cell, to more global patterns covering a few cells, allowing for subtle patterns to be captured even in the presence of stronger global patterns.

**Supplementary Fig. 3:**
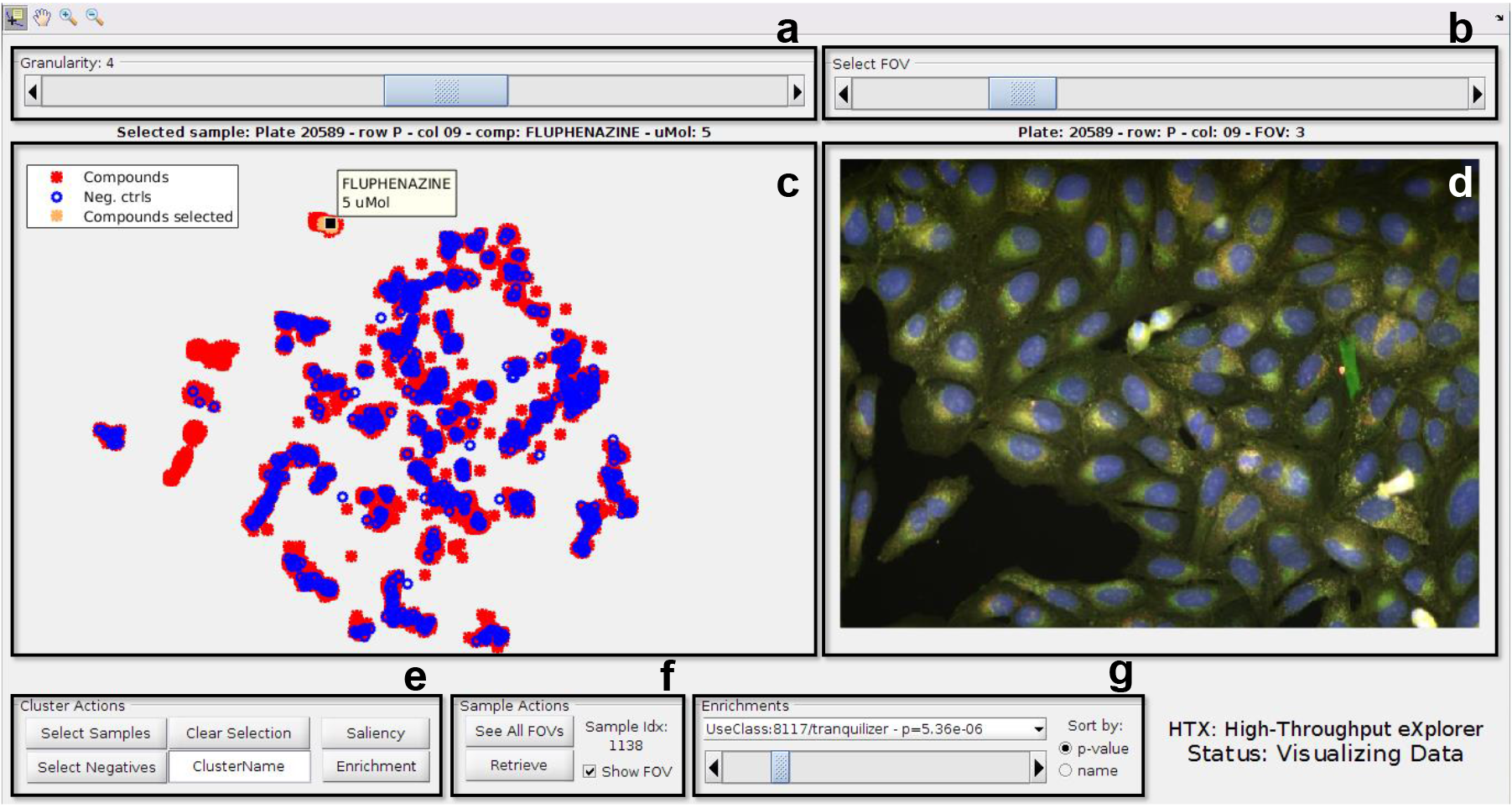
Description of the HTX demo interface. **(a)** the *granularity* control is used to navigate between the 2-D visualizations **(c)** at different levels of detail (see **Fig. 1a**). (b) the field-of-view (*FOV*) control is used to see the multiple images that belong to the same sample, which are plotted below **(d). (e)** the *cluster actions* panel contains the tools to select groups of samples from the 2-D maps, and perform the exploration actions of saliency and enrichment computation. By default, the saliency is computed against the control samples, but this can be changed by selecting “negative” samples from the visualization, as demonstrated in **Supplementary Video 1. (f)** when the cursor is selecting a sample in the 2-D map, the *cluster actions* panel offers an option to retrieve its top most similar samples. Finally, **(g)** the *enrichments* panel shows the enrichments pre-computed on the dataset using the available metadata, and allows the user to navigate through them by automatically selecting the sample neighbourhoods that were found to be enriched, as demonstrated in **Supplementary Video 2.**

